## Supplementary Figures for "Molecular dynamics of the matrisome across sea anemone life history"

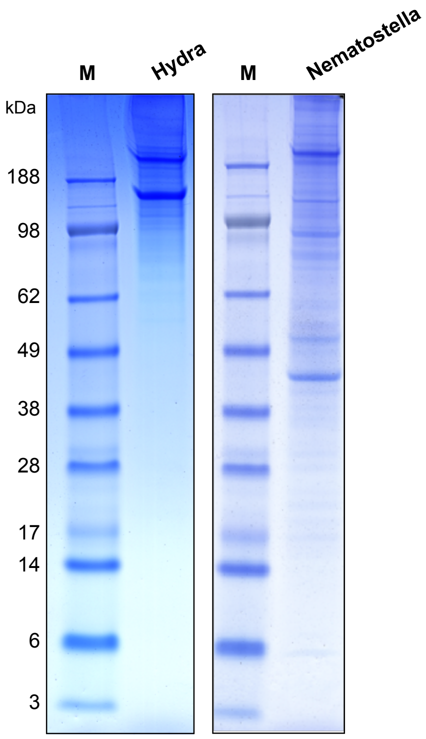


**Supplementary Figure S1. Protein complexity in mesoglea samples of *Nematostella* and *Hydra* resolved by one-dimensional SDS-PAGE.** Mesogleas were prepared from adult animals and dissolved in lithium dodecylsulfate buffer containing 1 M DTT. After heating at 90° C for 30 minutes, 130 mg of each sample was loaded on a 4-12 % gradient gel. The molecular mass of marker proteins (M) is as indicated.

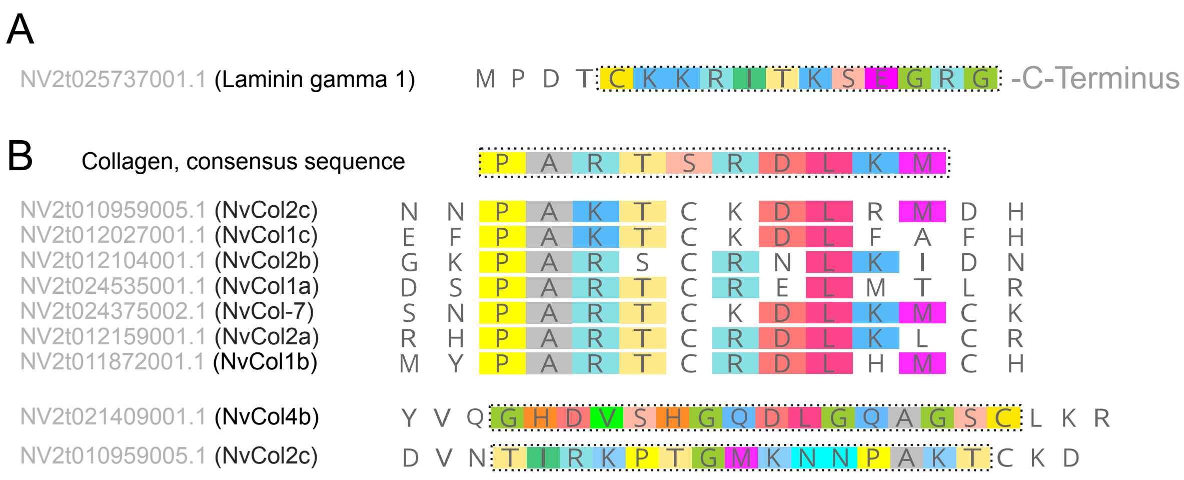


**Supplementary Figure S2. Epitope sequences of laminin and collagen antibodies.** (A) The laminin antibody was raised against a peptide sequence of the Nv laminin gamma 1 chain indicated by the boxed area. (B) The collagen antibodies were raised against an epitope representing a consensus sequence of fibrillar collagens (NvPanCol) and unique sequences in NvCol4b and NvCol2c as indicated. Mark that in the consensus sequence the central cysteine residue was replaced by a serine to prevent disulfide formation.

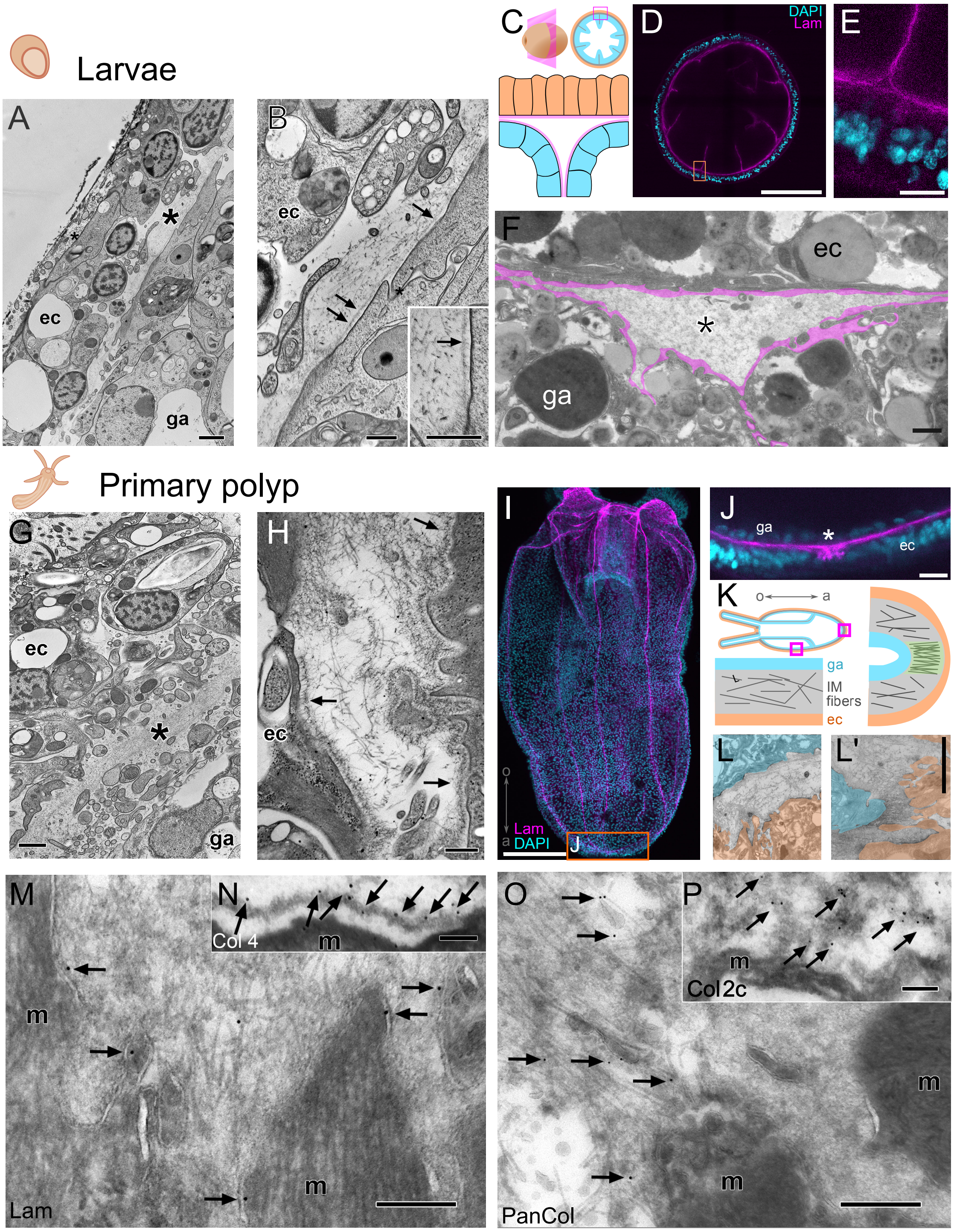


**Supplementary Figure S3. Ultrastructure of the mesoglea in *Nematostella* larvae (A-F) and primary polyps (G-P).** (A) Electron microscopy of the larvae with ectoderm (ec), gastroderm (ga) and an only 0.5 µm thick mesoglea (asterisk) in between. (B) The IM of cryofixed larval mesoglea displays scarce, quite thin filaments, embedded in amorphous compounds. The basement membrane (BM: arrows) appears rather delicate at this developmental stage, measuring about 70nm. (C) The mesoglea thickens into a triangular shape at the septa branching points. (D) Cross section of a larva stained with DAPI (cyan) and laminin antibody (magenta). DAPI hardly enters the gastroderm. The mesoglea and developing septae are shown by laminin staining. (E) The triangular expansion of the mesoglea at the septa branch is lined with laminin staining while the central region does not show any signal. (F) Electron micrograph of the septa branch. The BM is overlaid in magenta. The central region of the septa triangle is filled with loose fibrillar material. (Scale bars: A, F =1 µm, B = 0.5 µm, D = 100 µm, E = 10 µm). (G, H) EM of the primary polyp mesoglea (asterisk) and adjacent epithelia (ec, ga). The mesoglea is now about 1.5 µm thick with numerous, haphazardly arranged about 13nm thick and 6 nm thin fibrils in the IM. The BM (arrows) has thickened, thus, matured as well, and forms distinct, dense, about 130 nm thick meshworks lining the epithelia. (I) Whole mount staining of a tentacle bud stage stained with DAPI and laminin antibody. (J) At the aboral end, the BM is expanded (asterisk) into the ectodermal layer forming a knot-like structure. (K-L') Scheme and electron micrographs of the ECM fiber orientation at different sites of the primary polyps' body (ec highlighted in rose, ga in blue, ECM in grey/green). (L) The filaments/fibrils in the region of the body column are loosely oriented along the oral-aboral (o-a) axis following the overall orientation of the mesoglea. (L') At the aboral end, the fibrils are densely packed forming a plug-like structure parallel to the o-a axis (overlaid in green). Scale bars: G = 1 µm; H = 0.5 µm; I = 100 µm, J = 10 µm. L, L' = 1 µm. (M-P) Immunoelectron microscopy of mesoglea compounds, performed on thawed cryosections. (M) Laminin immunogold label (arrows) along the plasma membrane of the ECM lining muscle cells (m). (N) Col4 label located at the BM. (O) PanCol predominantly found throughout the IM. (P) Col2c label in the IM. Scale bars: M-P = 0.25 µm.

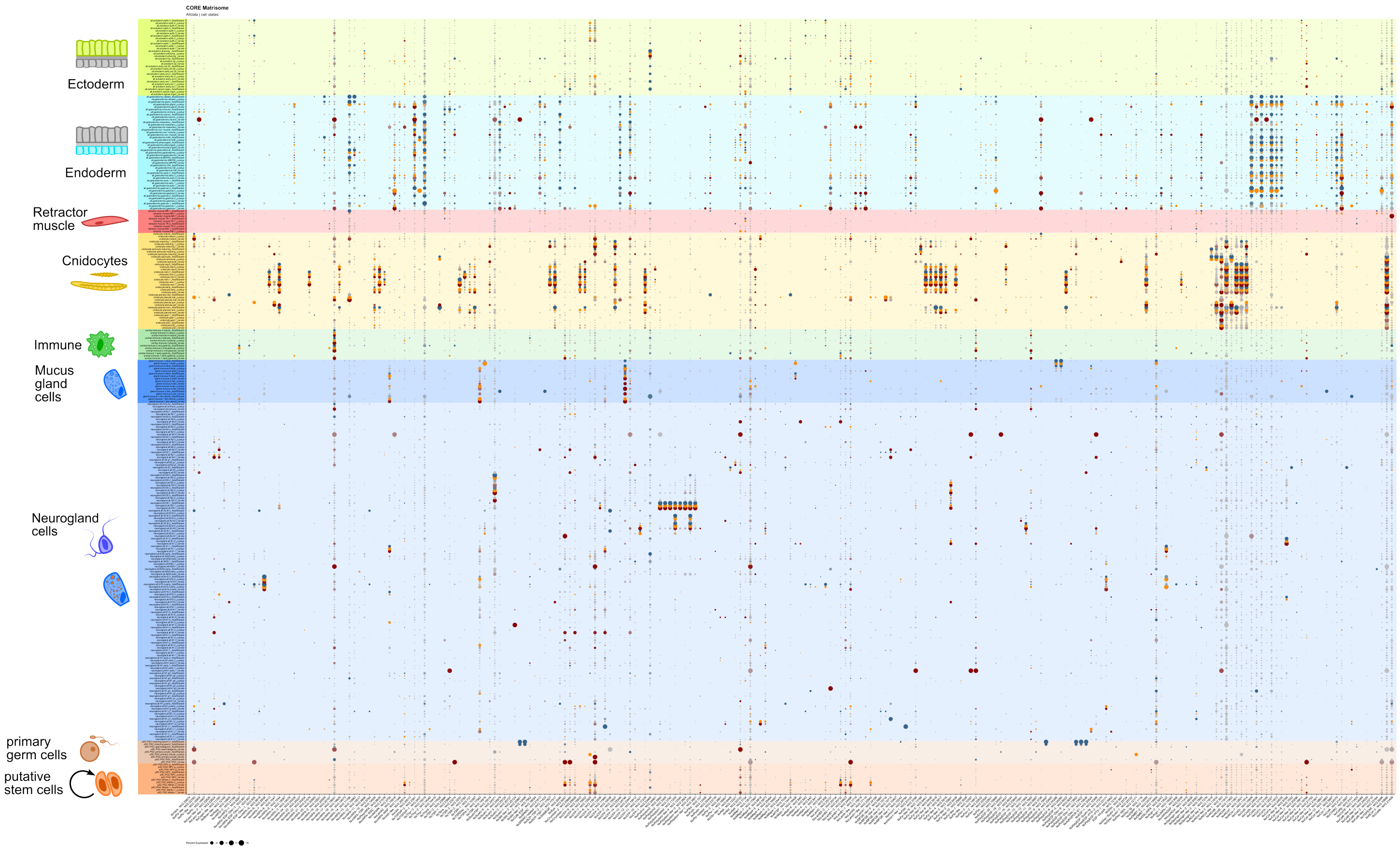


**Supplementary Figure S4. Expression profiles of CORE matrisome genes.** Dotplot expression profiles across all cell-state clusters, separated across phases of the life-cycle. Cell states are grouped and colored according to tissue-type partitions, and genes are grouped by category. Expression scale is the same as in Fig. 2. Abbreviations for cell state identities are explained in the README file on table S6.

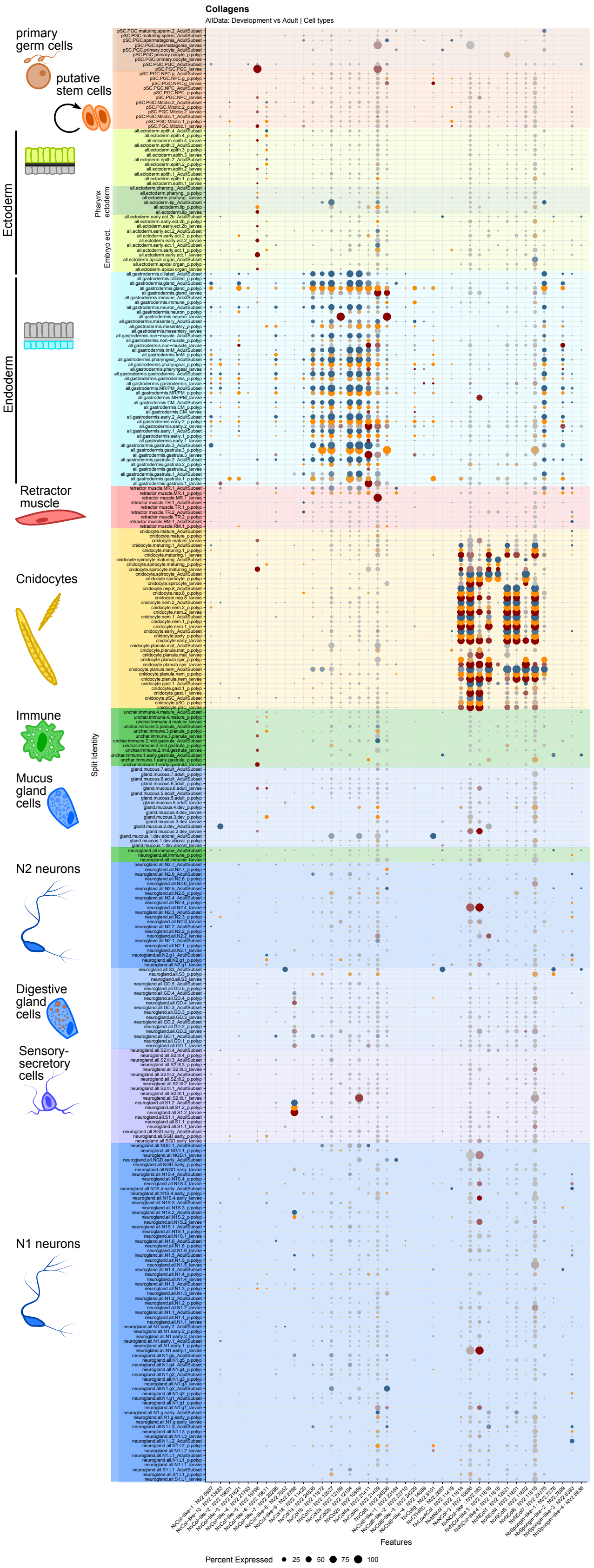


**Supplementary Figure S5. Single cell atlas of collagen genes.** Dotplot expression profiles of all identified collagen-coding genes across cell type partitions, separated across phases of the life-cycle. Cell states are grouped and colored according to tissue-type partition. Expression scale is the same as in Fig. 2. Abbreviations for cell identities are explained in the README file on Table S5.

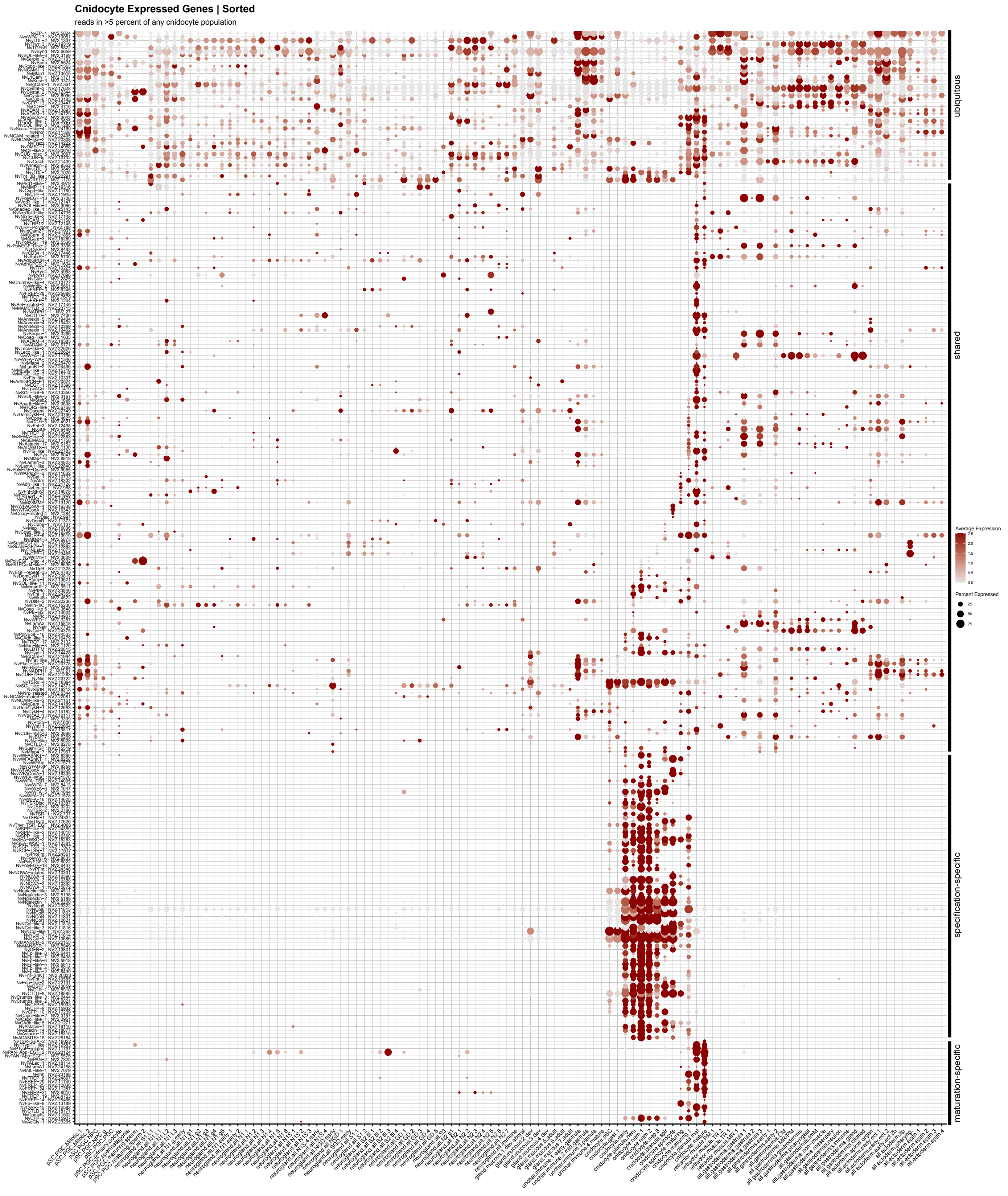


**Supplementary Figure S6. Single cell atlas of cnidocyte-expressed matrisome genes.** Dotplot expression profiles of matrisome genes expressed within the cnidocytes, plotted across all cell-type states. Genes are grouped according to degree of overlapping expression with other cell types, with ‘ubiquitous’ expression on top, followed by ‘shared’ expression, and then cnidocyte-specification specific genes, and mature cnidocyte-specific genes.

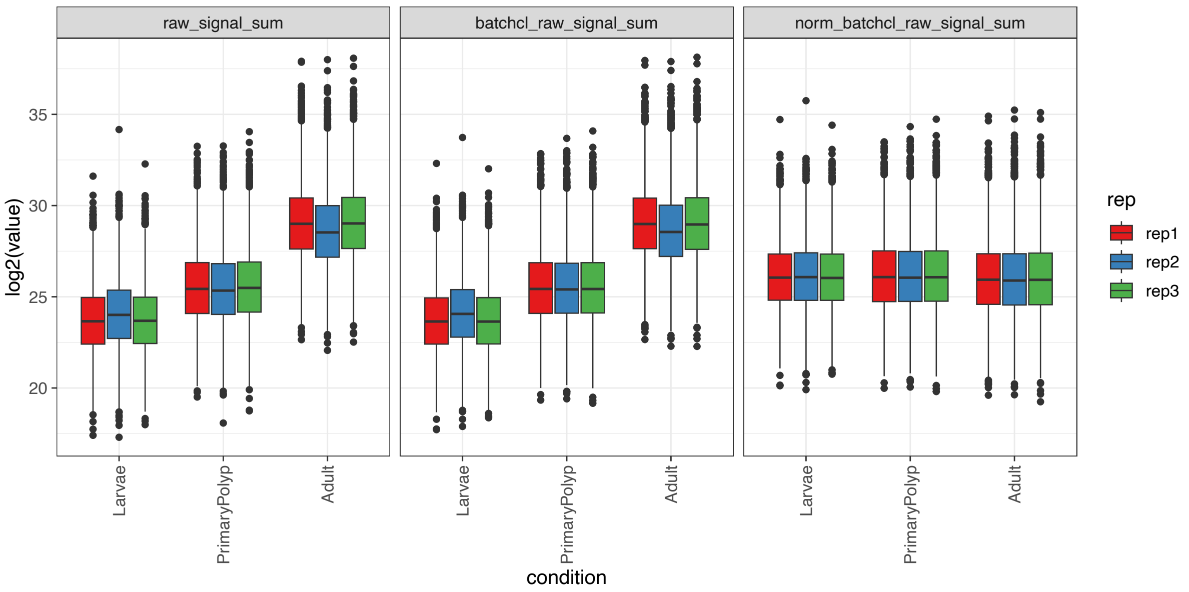


**Supplementary Figure S7. Mesoglea protein abundance overview in specific developmental stages.** Boxplot overview of data normalization steps for mesoglea samples. Raw TMT reporter ion intensities (left) were first cleaned for batch effects (middle) and further normalized using variance stabilization normalization (vsn - right). Due to low starting material in early cell stages, protein concentration was not adjusted before mass spec measurement and only accounted for in the normalization step to achieve equal protein amounts.

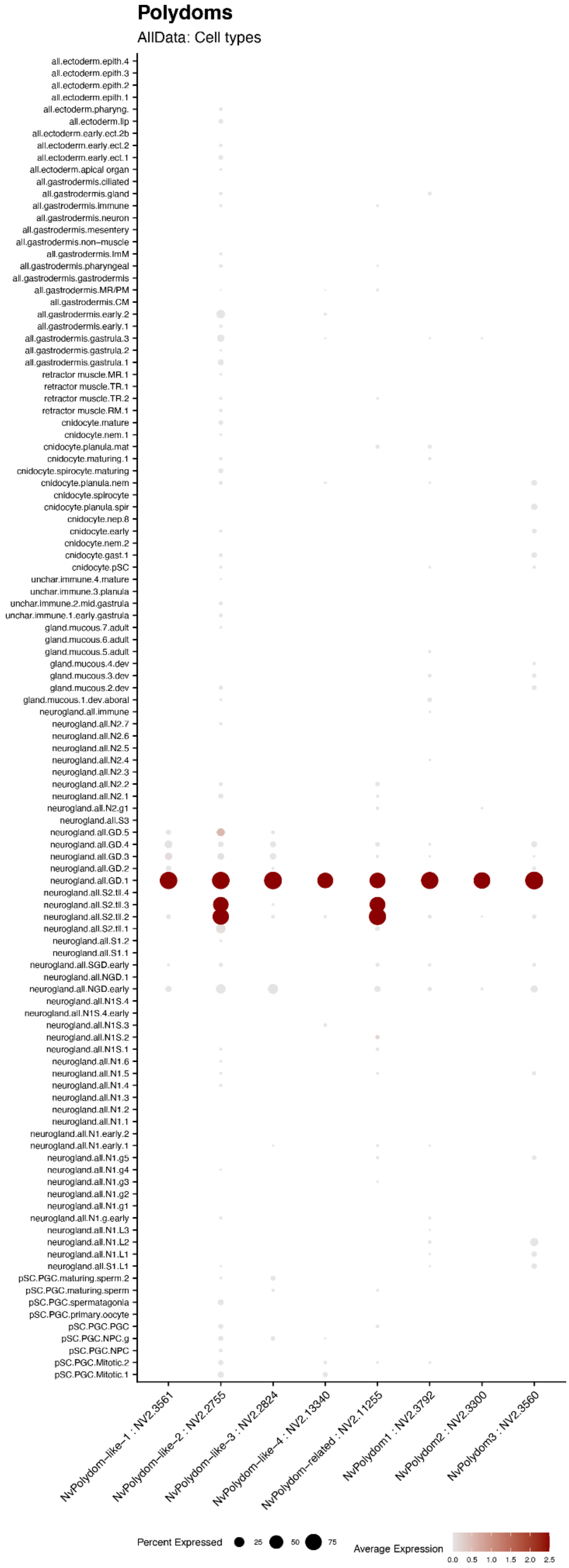


**Supplementary Figure S8. Expression profile of Nematostella polydoms.** All Nemastostella polydoms are expressed from GD.1 neurogland cells supposed to have a digestive function. Polydom-like-2 and Polydom-related show additional expression within the uncharacterised secretory cell type. S2.tII.2&3. function. Polydom-like-2 and Polydom-related show additional expression within the uncharacterised secretory cell type S2.tII.2&3.

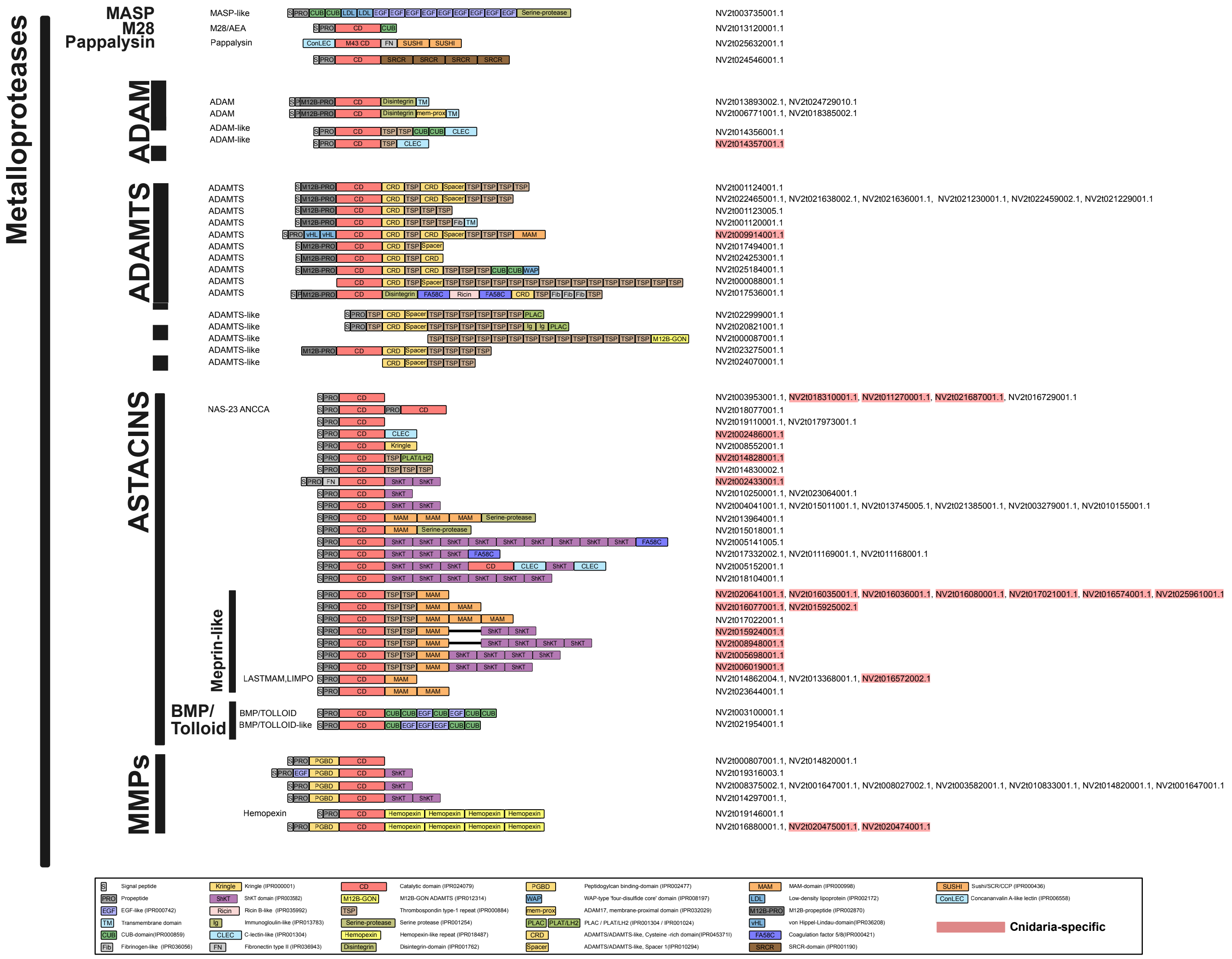


**Supplementary Figure S9. Domain organization of metalloprotases in the *Nematostella* matrisome.** Metalloproteases comprise an N-terminal signal-peptide for secretion (S), a propeptide or prodomain conferring latency (PRO) and a catalytic zinc-dependent metallopeptidase domain (CD). Additional domains were listed using Interproscan-5.57 codes: (Kringle) Kringle (IPR000001), (CD) Catalytic domain (IPR024079), (PGBD) Peptidogylcan binding - domain (IPR002477), (MAM) MAM-domain (IPR000998),(PRO) Propeptide, (ShKT) ShKT domain (IPR003582), (M12B-GON) M12B-GON-ADAMTS (IPR012314), (WAP) WAP-type 'four-disulfide core' domain (IPR008197), (LDL) Low-density lipoprotein (IPR002172), (EGF) EGF-like (IPR000742), (Ricin) Ricin B-like (IPR035992), (TSP) Thrombospondin type-1 repeat (IPR000884), (mem-prox) ADAM17, membrane-proximal domain (IPR032029), (M12B-PRO) M12B-propeptide (IPR002870), (TM) Transmembrane domain, (Ig) Immunoglobulin-like (IPR013783), (Serine-protease) Serine protease (IPR001254), (PLAC/PLAT/LH2) PLAC / PLAT/LH2 (IPR001304 / IPR001024), (vHL) von Hippel-Lindau-domain(IPR036208), (CUB) CUB-domain(IPR000859), (CLEC) C-lectin like (IPR001304), (Hemopexin) Hemopexin-like repeat (IPR018487), (CRD) ADAMTS/ADAMTS-like, Cysteine-rich domain(IPR045371I), (FA58C) Coagulation factor 5/8(IPR000421), (Fib) Fibrinogen-like (IPR036056), (FN) Fibronectin type II (IPR036943), (Disintegrin) Disintegrin-domain (IPR001762), and (Spacer) ADAMTS/ADAMTS-like, Spacer 1(IPR010294). All metalloproteases were identified based on manual sequence and domain analysis. The identified proteases were categorized as MASP-like (1), M28-like (1), ADAM (-like) (4), MMP (6), ADAMTS (-like) (15), and Astacin (27). The latter can be subdivided into Meprin-like (9), BMP/Tolloid-like (2) and other Astacins (16). Based on the identified hierarchical phylogenetic orthogroups we identified sequences that are specific to cnidarians as indicated.
