## Supplementary material for "Molecular dynamics of the matrisome across sea anemone life history": README File for Suppl. Tables

**README for Supplementary Tables**
====================

**Table S1**
*In silico matrisome prediction of the Nematostella mesoglea including results of differential abundance analysis using moderated t-test (limma) for each developmental comparison (Adult - Larvae, Larvae - Primary Polyp, Adult - Primary Polyp)*

transcript_id:
Stowers SIMRbase (https://simrbase.stowers.org/starletseaanemone) sequence ID from the Nematostella vectensis NV2 (wein_nvec200_tcsv2) transcriptome

StowersTranscriptSeqLink:
Link to the specific transcript record in the Stowers SIMRbase

gene_id:
Stowers gene ID without transcript version

Stowers Description:
Automatically generated description of the transcript by UniProt BLAST at the submission of the sequence to Stowers

new_manual_annotation:
Annotation from this publication based on literature research and sequence comparisons

short_name:
Suggested name also used in this publication

Division_manual:
Broad categories of the Adhesome: Core Matrisome, Matrisome-associated, or Adhesome

Categories_manual:
Subclasses of Adhesome proteins

Orthofinder_hierarchical_group:
Orthogroup number from evolutionary orthology search in this publication

Specificity:
Flag showing the evolutionary distribution of the gene based on the Orthofinder classification in our comparison, with a focus on Cnidaria, Anthozoan, and Nematostella specificity

closest_SwissProt_hit:
Closest BLAST hit against the SwissProt database at the time of publication

closest_SwissProt_description:
Description of the closest SwissProt BLAST hit at publication

fdr:
False discovery rate

hit_annotation:
Statistical classification of detected proteins as hit, candidate, or no hit

logFC:
Logarithmic fold change of protein in the comparison

pvalue:
MS detection p-value

**Table S2**
*Full mass spectrometry data of the Nematostella mesoglea*

qupm:
quantified unique peptide matches

top3:
average log10 MS1 intensity of the three most abundant peptides for a given protein

norm_batchcl_raw_signal_sum:
normalized and batch-cleaned reporter ion intensities from the TMT experiment

**Table S4**
Worksheet *MatrisomeExpression.cellstates*

Gene expression profiles for all matrisomal genes across all cell state identities in Cole et al. 2024, Frontiers in Zoology, https://doi.org/10.1186/s12983-024-00529-z.

Worksheet *MatrisomeExpression.lifecycle*

Gene expression profiles for all matrisomal genes across all cell state identities in Cole et al. 2024, Frontiers in Zoology, https://doi.org/10.1186/s12983-024-00529-z, with the cell states separated by life cycle stages used in the current paper: larva, primary polyp (p.polyp), and adult subsets.

avg.exp:
average expression

pct.exp:
the percentage of the cells with gene expression

avg.exp.scaled:
relative expression levels

cell.state.id:
cell state identities

pSC:
putative germ cell

PGC:
primary germ cell

NPC:
neuroglandular progenitor cell

GD:
digestive gland cell

NGD:
digestive neuroglandular cell

**Table S5**
*Single cell expression profiles of core matrisomal genes. Output of the FindAllMarkers function of the Seurat package (vs.4.4.0) run with default parameters using all core matrisome genes across all clusters annotated in Cole et al. 2024 Frontiers in Zoology, https://doi.org/10.1186/s12983-024-00529-z.*

cell.state.id:
cell state identities

gene:
core matrisome gene tested

p_val:
calculated p-value of the expression difference

avg_log2FC:
calculated average log2 fold change between cluster 1 (cell.state.id) and the rest of the dataset

pct.1:
percentage of cells in cluster 1 (cell.state.id) with gene expression

pct.2:
percentage of cells within the remaining clusters with gene expression

p_val_adj:
adjusted p-value

| Partition | ID | Full Name |
|---|---|---|
| pSC | | putative stem cells |
| | Mitotic.1 | pSC.mitotic number 1. putative S-phase mitotic cells. Express PCNA |
| | Mitotic.2 | pSC.mitotic number 2. putative M-phase mitotic cells. Expresses Nucleolar and spindle-associated protein 1 |
| | NPC | Neuroglandular precursor cells, express SoxC and SoxB(2) |
| | NPC.g | Neuroglandular precursor cells, express SoxC and Six1-2, gastrodermal |
| PGC | | Primary Germ Cells |
| | primary.oocyte | early primary oocytes |
| | spermatagonia | early spermatagonia |
| | maturing.sperm | maturing spermatagonia |
| | spermatozoa | spermatazoa |
| gland.mucous | | mucous gland, expresses mucin |
| | 1.dev.aboral | Mucin producing cell type, developmental cell state n.1, aborally enriched (expresses six3-6) |
| | 2.dev | Mucin producing cell type, developmental cell state n2 |
| | 3.dev | Mucin producing cell type, developmental cell state n3 |
| | 4.dev | Mucin producing cell type, developmental cell state n4 |
| | 5.adult | Mucin producing cell type, adult cell state n5 |
| | 6.adult | Mucin producing cell type, adult cell state n6 |
| | 7.adult | Mucin producing cell type, adult cell state n7 |
| unchar.immune | | uncharacterized cell state, putative immue response, expresses interferon regulatory factor (IRF1-2a) |
| | 1.early.gastrula | putative immune response, enriched in early gastrula |
| | 2.mid.gastrula | putative immune response, enriched in mid gastrula |
| | 3.planula | putative immune response, enriched in planula |
| | 4.mature | putative immune response, expresses interferon regulatory factor (IRF1-2a) |
| neuroglandular | | neuroglandular cells of secretory type |
| | S1.L1 | Secretory cell type 1, larval enriched number 1 |
| | N1.L1 | neuron N1 type, larval enriched number 1, expresses homeobox Nkx3 |
| | N1.L2 | neuron N1 type, larval enriched number 2, expresses PRGa, apical organ associated |
| | N1.L3 | neuron N1 type, larval enriched number 3 |
| | N1.g.early | neuron N1 type, inner cell layer derived, putative early cell state, expresses ashD |
| | N1.g1 | neuron N1 type, inner cell layer derived number 1, expresses homeodomain proteins OTX and Six1.2 |
| | N1.g2 | neuron N1 type, inner cell layer derived number 2, expresses homeodomain proteins OTX and Hox3-like |
| | N1.g3 | neuron N1 type, inner cell layer derived number 3, expresses retinal homeodomain protein (Rx) and pit1 |

| | | |
|---|---|---|
| | N1.g4 | neuron N1 type, inner cell layer derived number 4, expresses nkx2.2 |
| | N1.g5 | neuron N1 type, inner cell layer derived number 5, expresses pax2 |
| | N1.early.1 | neuron N1 type, putative early cell state number 1, expresses NK1 |
| | N1.early.2 | neuron N1 type, putative early cell state number 2, expresses transcription factor AP2 |
| | N1.1 | neuron N1 type number 1, expresses homeobox protein even-skipped homolog (evx) |
| | N1.2 | neuron N1 type number 2, expresses homeobox protein even-skipped homolog (evx) |
| | N1.3 | neuron N1 type number 3, expresses visual system homeobox protein (vsx) |
| | N1.4 | neuron N1 type number 4, expresses prdm13 |
| | N1.5 | neuron N1 type number 5, expresses nkx2.5 |
| | N1.6 | neuron N1 type number 6, putative early cell state, expresses GLWamide |
| | N1S.1 | sensory neuron N1 type number 1, expresses NOT homeobox |
| | N1S.2 | sensory neuron N1 type number 2, expresses gsx homeobox |
| | N1S.3 | sensory neuron N1 type number 3, expresses otp homeobox |
| | N1S.4.early | sensory neuron N1 type number 4, putative early state, expresses foxq2d2 homeobox |
| | N1S.4 | sensory neuron N1 type number 4, expresses foxq2d2 homeobox |
| | NGD.early | neurosecretory/gland putative early state, expresses homeodomain protein rough |
| | NGD.1 | neurosecretory/gland putative early state number 2, expresses homeodomain protein rough |
| | SGD.early | secretory/gland putative early state, expresses klf5 |
| | S1.1 | Secretory cell type, type1 number 1, expresses agrin |
| | S1.2 | Secretory cell type, type1 number 2, expresses DD3 |
| | S2.tll.1 | Secretory cell type, type2 number 1, expresses tll |
| | S2.tll.2 | Secretory cell type, type2 number 2, expresses tll |
| | S2.tll.3 | Secretory cell type, type2 number 3, expresses tll |
| | S2.tll.4 | Secretory cell type, type2 number 4, expresses tll |
| | GD.1 | Digestive gland cell type number 1 |
| | GD.2 | Digestive gland cell type number 2 |
| | GD.3 | Digestive gland cell type number 3 |
| | GD.4 | Digestive gland cell type number 4 |
| | GD.5 | Digestive gland cell type number 5 |
| | S3 | Secretory cell type 3 |
| | N2.g1 | gastrodermal neuron N2 type number 1, expresses prdm14d |

| | | |
|---|---|---|
| | N2.1 | neuron N2 type, putative early state, expresses fez/barh2 |
| | N2.2 | neuron N2 type, number 2, expresses QGRFa |
| | N2.3 | neuron N2 type, number 3, expresses fez |
| | N2.4 | neuron N2 type, number 4, expresses homeodomain protein NOT2 |
| | N2.5 | neuron N2 type, number 5, expresses homeodomain protein runx |
| | N2.6 | neuron N2 type, number 6, putative tentatcle retractor muscle early state1, expresses homeodomain protein Nkx2.2c, tbx20b, and nem64 |
| | N2.7 | neuron N2 type, number 7, putative tentatcle retractor muscle early state2, expresses homeodomain protein tbx20a, and nem64 |
| | immune | immune cell state, expresses NFKB1 |
| cnidocyte | | early cnidoctyes: undergoing capsule specification |
| | pSC | putative stem cell state |
| | gast.1 | gastrula-enriched cnidocytes |
| | planula.nem | planula-enriched nematocyte |
| | planula.spir | planula-enriched spirocyte |
| | planula.mat | planula-enriched mature state |
| | early | early cnidocyte cell state |
| | nem.1 | cnidocyte specification type nem.1 |
| | nem.2 | cnidocyte specification type nem.2 |
| | nep.8 | cnidocyte specification type nep.8 |
| | spirocyte | spirocyte specification state |
| | spirocyte.maturing | spirocyte mature state |
| | maturing.1 | cnidocyte maturation state |
| | mature | mature cnidocyte state |
| epithelia.ect | | outer epithelia |
| | apical organ | ectodermal cells of the apical organ |
| | early.ect.1 | early ectoderm state n1 |
| | early.ect.2 | early ectoderm state n2 |
| | early.ect.2b | early ectoderm state n1 |
| | lip | boundary ectoderm, gastrula enriched expresses FoxA |
| | pharyng. | outer epithelia of pharynx and septal filaments, expresses FoxA |
| | epith.1 | epithelial ectoderm cell state n1 |
| | epith.2 | epithelial ectoderm cell state n2 |
| | epith.3 | epithelial ectoderm cell state n3 |
| | epith.4 | epithelial ectoderm cell state n4 |
| retractor muscle | | retractor muscle: fast contracting type |
| | RM.1 | Retractor mucle mature cell state |
| | TR.2 | Tentacle retractor muscle cell state n2, expresses acetylcholine receptor and nem64 |

| | | |
|---|---|---|
| | TR.1 | Tentacle retractor muscle cell state n1, expresses wnt1 and nem64 |
| | MR.1 | Mesentary retractor muscle cell state n1, nem26 |
| gastrodermis | | gastrodermis, inner cell layer |
| | 1 | mature gastrodermis cell state n1 |
| | 2 | mature gastrodermis cell state n2 |
| | SlowMuscle | slow muscle: parietal and circular ring muscle |
| | RM&ImM | Retractor muscle and Intermuscular membrane |
| | pharyngeal | pharyngeal gastrodermis |
| Partition | ID | Full Name |
| pSC | | putative stem cells |
| | Mitotic.3 | pSC.mitotic number 1. putative S-phase mitotic cells. Express PCNA |
| | Mitotic.4 | pSC.mitotic number 2. putative M-phase mitotic cells. Expresses Nucleolar and spindle-associated protein 2 |
| | NPC | Neuroglandular precursor cells, express SoxC and SoxB(2) |
| | NPC.g | Neuroglandular precursor cells, express SoxC and Six1-2, gastrodermal |
| PGC | | Primary Germ Cells |
| | primary.oocyte | early primary oocytes |
| | spermatagonia | early spermatagonia |
| | maturing.sperm | maturing spermatagonia |
| | spermatozoa | spermatazoa |
| gland.mucous | | mucous gland, expresses mucin |
| | 1.dev.aboral | Mucin producing cell type, developmental cell state n.1, aborally enriched (expresses six3-6) |

**Table S6**

*Matrix of average expression values for all cnidocyte-specific genes (rows) across all cell state identities in Cole et al. 2024 Frontiers in Zoology, [https://doi.org/10.1186/s12983-024-00529-z](https://doi.org/10.1186/s12983-024-00529-z).* Cell IDs are as in Table S5.

**Table S7**

*Quantitative mass spectrometry data of mesoglea samples from different life stages.*

logFC:
Logarithmic fold change of protein in the comparison

-log10(pvalue):
MS detection p-value
